# Heritable and environmental influences on innate immunity across three populations of threespine stickleback (*Gasterosteus aculeatus*)

**DOI:** 10.1101/2025.07.11.664444

**Authors:** Cole J. Wolf, Panna A. Codner, Jesse N. Weber

**Affiliations:** Department of Integrative Biology, University of Wisconsin–Madison, Madison, WI 53706, USA

**Keywords:** innate immunity, ecoimmunology, environmental plasticity, oxidative burst

## Abstract

Environmental variation plays a key role in immune development and function; factors such as pathogen exposure history, seasonality, and resource availability all affect an individual’s immune phenotype. However, the relative contributions of heritable and non-heritable factors remain unclear for most immune phenotypes. We used three populations of threespine stickleback (*Gasterosteus aculeatus*) with heritable differences in immune function to investigate the relationship between immunity, genetic divergence, and the environment. To test for environmental effects on immunity, fish were raised in tanks with different flow rates (continuous or intermittent). After long-term acclimation to one tank environment, subsets of adult fish were moved to the alternate flow regime and allowed to acclimate for eighteen weeks. We then measured the effects of starting environment, transfer between environments, and final environment across several immune parameters. Fish population and treatment both significantly affected immune function. Stickleback from a population previously found to display the highest heritable levels of innate immunity displayed the highest oxidative burst capacity (ROS) regardless of tank environment. However, all fish in intermittent flow tanks (both resident and transfers) tended to have higher ROS production, more granulocytes, and greater spleen mass than in continuous flow. Variation in liver mass was mainly driven by population effects. We also provide limited data suggesting that the two water flow regimes harbor different microbial environments, offering a potential future direction for understanding the proximate connection between tank environment and immune variation. Overall, this work demonstrates that a simple change in water flow dynamics can induce immune flexibility. Perhaps more importantly, it also highlights the need for further research examining how naturally evolved genetic differences and environmental factors individually and jointly influence the magnitude and direction of immune responses.

## Introduction

Differential immune responses facilitate an organism’s ability to maintain high fitness in the face of changing pathogen risk. While some elements of immunity are highly innate, others are influenced primarily by either environment or by genotype-by-environment interactions. Ecoimmunology has traditionally focused on environmental factors such as seasonality, resource availability, and exposure to pathogens (Schoenle et al. 2018). This is largely due to difficulty in identifying heritable differences in wild organisms, which is further complicated when environmental influences vary across the life of an organism. For example, many species have a critical window where immune system development is sensitive to external inputs. In these cases, the adult immune function is contingent on early life exposures (MacGillivray and Kollmann 2014; Stinson 2020). In contrast, other aspects of immunity display plastic shifts depending on current environmental conditions. This includes the large body of work examining how exposure to toxins impacts immunity and disease (e.g., Bols et al. 2001; Salo et al. 2005; Lage et al. 2006; Coors et al. 2008; Kreitinger et al. 2016; Melchiorre et al. 2023). However, less is known about whether environmentally induced immune shifts are common to all individuals of a species or parceled into genotype-specific effects where only some individuals respond to cues (i.e., genotype-environment interactions; Bolnick et al. 2025).

The microbial environment surrounding an organism has strong impacts on host immunity. In mammals, the gut microbiome plays an important role in immune development and function, including maintenance of homeostasis and disease resistance (Zheng et al. 2020). Much of what we know in this area comes from laboratory work on captive, inbred lines of house mice (*Mus musculus*). In addition to their limited genetic diversity, housing mice in lab environments may limit our ability to establish links between microbes and host immune phenotypes. Indeed, lab-reared mice lack certain immune cell populations that are found in free-living mice, but these cell populations develop after co-housing with unrelated pet store mice (Beura et al. 2016). Laboratory mice also display much lower levels of immune activation than wild animals, likely due to their limited pathogen exposure (Abolins et al. 2017). Wild immune phenotypes can also be recapitulated through “rewilding” experiments that acclimate lab mice to natural outdoor enclosures. For example, wild-acclimated mice greatly increase granulocyte production and immune activation relative to laboratory animals (Lin et al. 2020; Yeung et al. 2020). Similarly, inoculating lab mice with wild gut microbiomes lowered levels of inflammation and improved resistance to viral infection (Rosshart et al. 2017). Although most immunology research has focused on rodent models, teleost fish represent an emerging model for connecting environmental and microbial variation with immune development and function (Jimoh et al. 2025).

Fish models offer several useful avenues for studying immune variation. One clear benefit of these systems is that their mucosal organs are in constant contact with the aquatic environment and can quickly respond to environmental changes (Jimoh et al. 2025). Examining not only the unique immune functions of fish mucosal tissues, but also their response to environmental variation, is an area of active research (Smith et al. 2026). A substantial amount of fish research has also addressed connections between pathogens and innate immunity, mostly focusing on organ specific immune responses (reviewed in Sayyaf Dezfuli et al. 2023). Apart from tissue or response specific studies, the ability to easily breed and rear fish across a spectrum of environments also offers research opportunities that are often not available with rodent models. For example, comparing similar strains housed in either lab, controlled outdoor (i.e., aquaculture), or wild settings offers opportunities to disentangle genetic and environmental effects on immunity (Zelikoff et al. 2000; Robertson et al. 2016). Moreover, recent work has identified several fish species harboring genetic variation with large effects on immune traits (Peuß et al. 2020; Weber et al. 2022), opening opportunities to further explore interactions between genes and environment.

Threespine stickleback fish (*Gasterosteus aculeatus*) are well known as models for studying the process of adaptation via natural selection. This ancestrally marine species repeatedly colonized and adapted to freshwater ecosystems across the northern hemisphere as glaciers receded 11,000 years ago (Bell and Foster 1994; McKinnon and Rundle 2002; Reid et al. 2021). Recent work suggests that population-specific immune divergence is an important factor in the adaptive radiation of stickleback into freshwater environments. Differences in innate immune function have been documented between numerous ecotypes, including marine-freshwater (Whiting et al. 2018), sympatric benthic-limnetic (Stutz et al. 2014), and parapatric lake-stream populations (Scharsack et al. 2007), as well as over larger geographic scales (Hamley et al. 2017; Weber, Kalbe, et al. 2017; Piecyk et al. 2021). Many of these studies rely on two classic assays of immune cells collected from the head kidney (i.e., the primary hematopoietic organ in fish), often involving flow cytometric methods (e.g., Weber, Steinel, et al. 2017; Weber et al. 2022). In one case, forward and side scatter are used to categorize populations and relative ratios of granulocytes and lymphocytes in the head kidney, with greater proportions of granulocytes suggesting greater investment in innate immunity. The second assay measures the oxidative burst response of granulocytes, where fluorescent dyes allow the quantification of reactive oxygen species (ROS) production after cells are exposed to an immune stimulant. In general, populations with high natural macroparasite exposure tend to evolve higher proportions of granulocytes and larger oxidative burst responses. Apart from flow cytometric assays, splenosomatic index (spleen mass/body mass, SSI) is frequently used to evaluate immune activity in fish, with an increased SSI corresponding to higher pathogen related immune response (Kaufmann et al. 2017; Rajkov et al. 2021), potentially resulting from a greater need to filter circulating leukocytes. Similarly, hepatosomatic index (liver mass/body mass, HSI) provides a proxy for individual condition and energy reserves (Chellappa et al. 1995), which may influence immune competency.

We took advantage of threespine stickleback populations with recently evolved heritable differences in immune function to examine the effects of both genetics and environment on immune variation. Specifically, we reared three populations of fish in a controlled laboratory setting and assayed their immune responses after exposure to two water flow environments (i.e., constant versus intermittent flow). This included contrasts between long-term rearing in one environment versus transfers to a new environment. The immune responses of two populations used in this study have been characterized previously (Weber, Steinel, et al. 2017; Weber et al. 2022). Fish from Roberts Lake, a tapeworm-resistant freshwater population on Vancouver Island, exhibited high oxidative burst responses. In contrast, fish from nearby Gosling Lake displayed a tapeworm-tolerant phenotype with relatively low oxidative burst capacity. However, the Gosling Lake population recently experienced a large introgression event resulting in a large genomic turnover, including changes in traits (i.e., peritoneal fibrosis) and genes known to be associated with tapeworm immunity (Flanagan et al. 2025). Our experiment sampled Gosling Lake fish after the introgression and, to our knowledge, the cellular immune responses of these fish have not been extensively measured post introgression. The third population (Kenai River Estuary in the Kenai Peninsula) represents a marine ecotype, the ancestral form of threespine stickleback (McKinnon and Rundle 2002), with unknown levels of ROS-associated immune responses. A previous study also reported that fish in intermittent flow tanks had higher oxidative burst responses than animals housed in constant flow conditions (Weber et al. 2022), so we expected to see the same pattern in this experiment. We also predicted that adults would display environment-specific flexibility; that is, the level of immune response would primarily be associated with current (not previous) water conditions. Finally, we hypothesized that variation in water flow rate would affect the composition of microbes in our fish tanks. We predicted that intermittent flow tanks would have more diverse microbial communities due to higher environmental stability and the presence of a sponge biofilter in each tank, and that this diversity may explain previous reports of higher oxidate burst levels in intermittent flow tanks. Although most of our measurements focused on cellular immunity, to understand more generalized effects of water flow we also compared plasticity in two organs. We predicted that increased immune investment would correlate with increases in spleen size, and that decreased liver size would indicate increased energetic expenditure, potentially connected to immune activation.

## Materials and Methods

### Animal Care & Husbandry

Wild adult fish (F0-generation) were sampled from two populations on Vancouver Island, British Columbia, Canada: Roberts Lake (Lat: 50.222, Long: -125.541) and Gosling Lake (Lat: 50.061, Long: -125.504). Details of the heritable immune differences observed between these populations has been reported previously (Weber, Steinel, et al. 2017; Weber et al. 2022). The other population consisted of marine fish (hereafter “marine”) from the Kenai River Estuary in Alaska, USA (Lat: 60.542, Long: -151.218). Multiple within population crosses were made via in-vitro fertilization in the field in summer 2019 to create an F1-generation. These fertilized eggs were transported to the University of Alaska-Anchorage (ADFG license P-19-005; Canada: NA19-457335). After reaching maturity in 2021, adult F1-fish were then transported to an aquarium at the University of Wisconsin-Madison. After this last transfer, fish were bred with unrelated individuals of the same population in the lab, and those offspring (F2-generation) were used in this experiment. This breeding design should minimize maternal effects on measured phenotypes, as all F1-parents shared the same laboratory environments. After hatching and prior to experimental manipulations, fish were housed in 18L tanks in full-sibling groups (initial n/tank = 2 – 15) with a water temperature of 17°C. The room was kept on a 16h light:8h dark cycle. Fish were fed daily with a mixture of freeze-dried bloodworms and mysis shrimp, and each tank had a section of PVC pipe and two artificial plants for environmental enrichment. All husbandry and experimental protocols were approved by the Institutional Animal Care and Use Committees at the University of Alaska-Anchorage (protocol #: 1308953-2) and the University of Wisconsin-Madison (protocol #: L006460).

### Acclimation Design

We used two water flow regimes to create different tank environments. Continuous-flow tanks had a volume of 18L and input flow rate of 1L/minute. The intermittent flow tanks had a volume of 23L, and water changes involved 1L/minute for 5 minutes every 12 hours. The intermittent flow tanks were also outfitted with bubbling sponge biofilters (Bioneat, small size) to maintain water quality and provide oxygenation. All tanks in the experiment shared a common water system, and water was run through a UV filter before recirculation to limit microbe movement between tanks.

As noted previously, after hatching all fish were initially reared in constant flow tanks with siblings, but approximately half of the tanks were then housed for at least 11 months in intermittent flow tanks. When fish were between 14 – 24 months in age, each family was randomly assigned into one of four treatments: continued exposure to continuous water flow (‘c’), continued exposure to intermittent flow (‘i’), transition from continuous to intermittent flow (‘ci’), and transition from intermittent to continuous flow (‘ic’). Both transition treatments involved moving families into new tanks, while fish in the continued exposure treatments were left in their original tanks (sample size and family structure is described in Supplemental Table 1). After approximately eighteen weeks in these experimental conditions, we fasted fish for 24 hours and then performed euthanasia via a combination of MS-222 overdose (500 mg/L) and pithing. We recorded each individual’s mass and standard length, then performed dissections to remove and weigh the liver and spleen. Head kidneys (HKs) were also collected for flow cytometric assays.

### Flow Cytometry Assay

Maximum ROS production in HK cells was quantified via oxidative burst assays using a previously described protocol (Weber, Steinel, et al. 2017; Weber et al. 2022). Briefly, HKs were initially placed in ice cold media (0.9× Roswell Park Memorial Institute medium containing 10% fetal bovine serum, 100 μM nonessential amino acids, 100 U/mL penicillin, 100 μg/mL streptomycin, and 55 μM β-mercaptoethanol). Single cell suspensions were generated by manually disrupting and filtering the tissue on a cell strainer (35 μm) with a pipette. We washed the cell suspension once in 4 mL of cold media, pelleted cells via centrifugation (300 xg, 4°C, 10 min), poured off most of the supernatant, and then resuspended the cell pellet in the remaining media. Live HK cells were counted using a hemocytometer (Hausser Scientific 3520) via Trypan blue exclusion (Corning 25-900-CI). We then distributed 2×10^5 HK cells into three separate wells on a 96-well round bottom plate. The first well also received 200 μL of HK medium, while the second and third wells received HK media with DHR-123 (ROS indicator fluorochrome, 2 μg/mL, Sigma D1054). Samples were then incubated at 18°C, 3% CO_2_ for 10 minutes. Next, 15 μL of PMA (Phorbol 12-myristate-13-acetat, 130 ng/mL; Sigma P8139) was added to the third well to stimulate ROS production, while wells 1 and 2 received equivalent volumes of HK media. Cells were incubated for a final 20 min at 18°C, 3% CO_2_, placed in a sealed bag on ice for transport, and assayed on a ThermoFisher Attune NxT V6 flow cytometer.

### Data Analysis

Flow cytometry data was analyzed in FlowJo (TreeStar). The magnitude of ROS production during oxidative burst assays was determined by comparing median fluorescence intensity of PMA-stimulated and unstimulated cells. The proportion of granulocytes and lymphocytes was quantified in the unstained, unstimulated HK cells using forward and side scatter gating (Weber, Steinel, et al. 2017; Weber et al. 2022).

All other data analysis was done in R (v4.2; R Core Team 2025). Because the amount of ROS produced by head kidney cells depended on time since exposure to PMA, we included flow cytometry sample order as a covariate. Specifically, we made separate linear models for PMA-stimulated and unstimulated cells with sample order as a random effect (R package *lme4* v35.5; Bates et al. 2015). We extracted the model residuals to represent time-corrected median fluorescence intensity and then subtracted the unstimulated value from the PMA-stimulated cells to control for individual variation in background ROS production (hereafter ROS residMFI), which we used as the response variable for all models of ROS production. We then built a series of mixed effect linear models that included one of four response variables (ROS residMFI, percent granulocytes, or mass of either liver or spleen) and a suite of potential predictor variables. We included fixed effects of population, sex, flow treatment (c/ci/ic/i), final flow environment (continuous/intermittent), movement between tanks (fish in the ic and ci groups were moved, c and i were not), body mass (only used for spleen and liver), and all combinations of predictor variables and pairwise interaction terms between population and one predictor. All models included fish family and age in days as additive random effects. We used the R package *performance* v12.4 (Lüdecke et al. 2021) to compare models and selected the model with the lowest AIC score for each response variable. The R package *emmeans* v1.11 (Lenth and Piaskowski 2026) was used to estimate group averages within a predictor variable, and to calculate Cohen’s effect size (d) to test for significant differences between groups.

### Tank 16S Sequencing & Analysis

We performed DNAsequencing on a subset of tanks to characterize microbial community differences between flow regimes. Five continuous and five intermittent flow tanks were sampled using a microbiome testing kit from Aquabiomics LLC (Junction City, OR, USA). Two samples were taken from each tank. First, 60 mL of tank water was slowly pushed through a 22-micron mixed cellulose ester filter, followed by 2 mL of 90% ethanol; the filter was allowed to dry then sealed in a small plastic container. Second, each tank’s water intake tube was swabbed vigorously with a cotton swab. The swab was then dunked in a 1.5 mL cryotube filled with 90% ethanol, allowed to dry, and sealed in a plastic tube. Samples were then shipped to Aquabiomics, which extracted DNA with Omega Bio-Tek Mag-Bind® kits, used universal primers to amplify the variable region of the 16S ribosomal RNA, and sequenced resulting amplicons using 2x500bp reads on an Illumina MiSeq.

Sequence data were curated with the *DADA2* pipeline (v1.16; Callahan et al. 2016). Reads were filtered and trimmed (command: ‘filterAndTrim’) with parameters maxN=0, truncQ=2, rm.phix=TRUE and maxEE=c(2,5). Paired reads were then merged (command: ‘mergePairs’) and chimera sequences were removed (command: ‘removeBimeraDenovo’). R package *DECIPHER* (v.2.26; Murali et al. 2018) was used to construct an amplicon sequence variants (ASVs) table (command: ‘IdTaxa’) after training on a reference dataset (SILVA 138 SSU; Quast et al. 2013). Analyses of species richness and differences between communities were completed with the R package *phyloseq* (v1.42; McMurdie and Holmes 2013).

## Results

### Oxidative burst assays

The best fit regression model for ROS residMFI during oxidative burst assays included fixed effects for population and flow treatment, and age and family as random effects. Across all treatments, the Roberts Lake stickleback produced significantly higher ROS than Gosling Lake (mean of 0.55 residMFI vs -0.48 residMFI, Cohen’s d =3.56, t-test p<0.00001, Figure 1) and marine populations (d =1.72, p<0.00001). The Gosling Lake population also produced less ROS than marine fish (d=-1.84, p<0.00001). These data support a strong role of population ancestry and evolutionary divergence in shaping oxidative burst capacity. After accounting for population effects, the c treatment had the lowest ROS production, which was significantly lower than the i (d=-0.27, p = 0.004) and ci treatments (d=-0.505, p = 0.0001) and marginally lower than the ic treatment (d=0.37, p = 0.051). Movement between flow environments (i.e., treated as a binary trait) was associated with increased ROS production but not included in the top model. Similarly, fish whose final environment was intermittent flow tanks tended to have greater ROS production than those in continuous flow (d=-0.11, p = 0.002), but including this variable did not improve the model fit after accounting for four-way treatment category. There was no effect of sex on ROS generation (p = 0.74). There was a positive correlation between ROS residMFI and fish age (Pearson’s correlation coefficient=0.55, p<0.00001)

**Figure 1.**
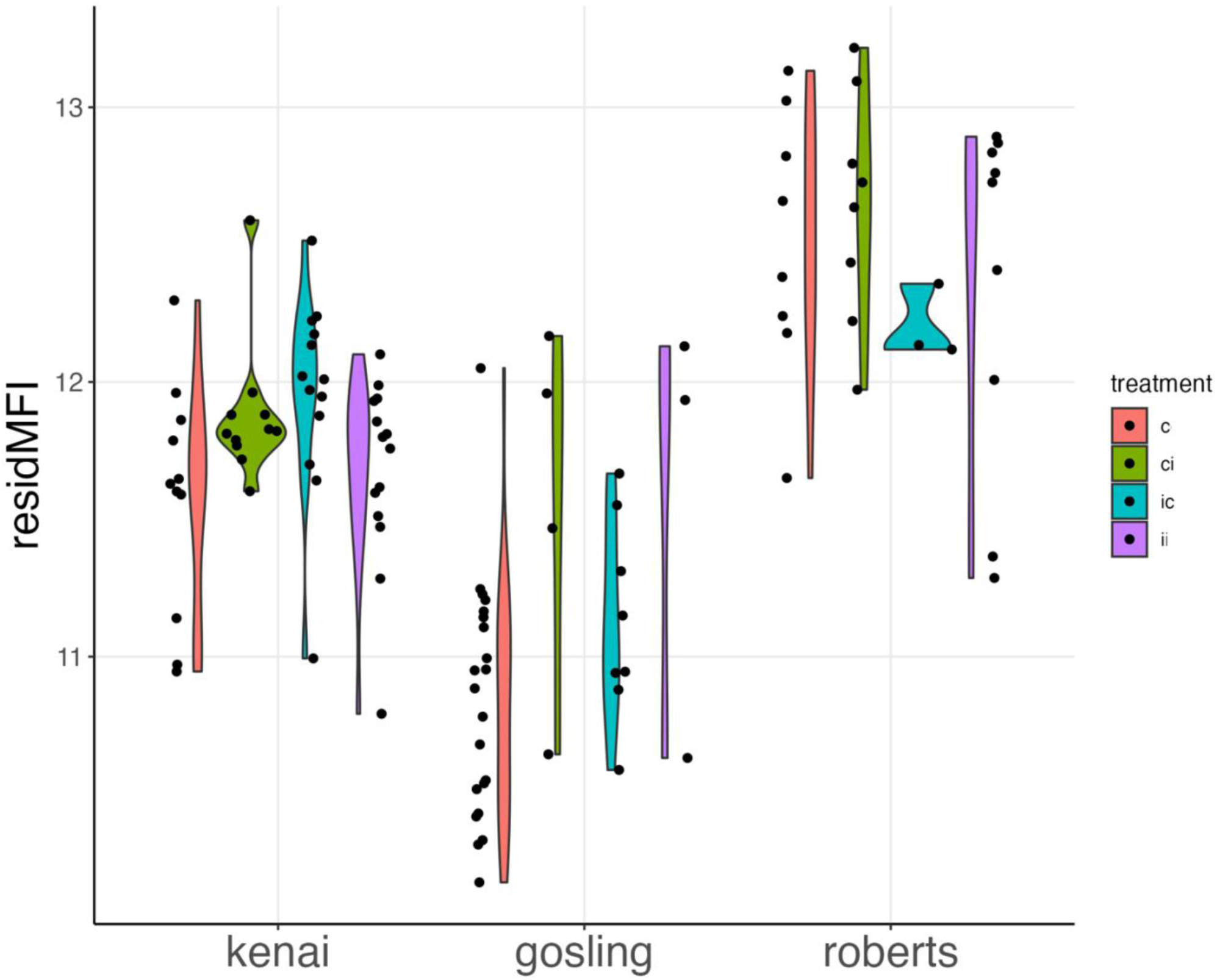
ROS production (residual MFI) during oxidative burst assay in head kidney cells by population. Color indicates experimental treatment: continuous only (“c” = red), continuous to intermittent (“ci” = green), intermittent to continuous (“ic” = blue), and intermittent only (“i” = purple).

### Granulocyte:Lymphocyte Ratio

The best supported model for percent granulocytes included additive effects for population, sex, and final flow environment, and age and family as random effects. Four-way comparisons of flow treatment and movement between flow environments were not associated with significant changes in percent granulocytes. The proportion of granulocytes in head kidneys of fish from intermittent flow tanks (ci and i treatments combined) was significantly than those removed from continuous flow tanks (38.1% vs 34%; t-test t = -3.08, p = 0.003, Cohen’s d =-0.03, Figure 2). Fish from Roberts Lake also had higher proportions of granulocytes than the Gosling Lake (41.2% vs 34.4%, p = 0.02, d=0.83) or marine (32.6%, p = 0.001, d=1.05) populations. The latter two populations were not significantly different (p = 0.47). There was a positive correlation between fish age and granulocyte proportion (Pearson’s correlation coefficient=0.32, p=0.0002)

**Figure 2.**
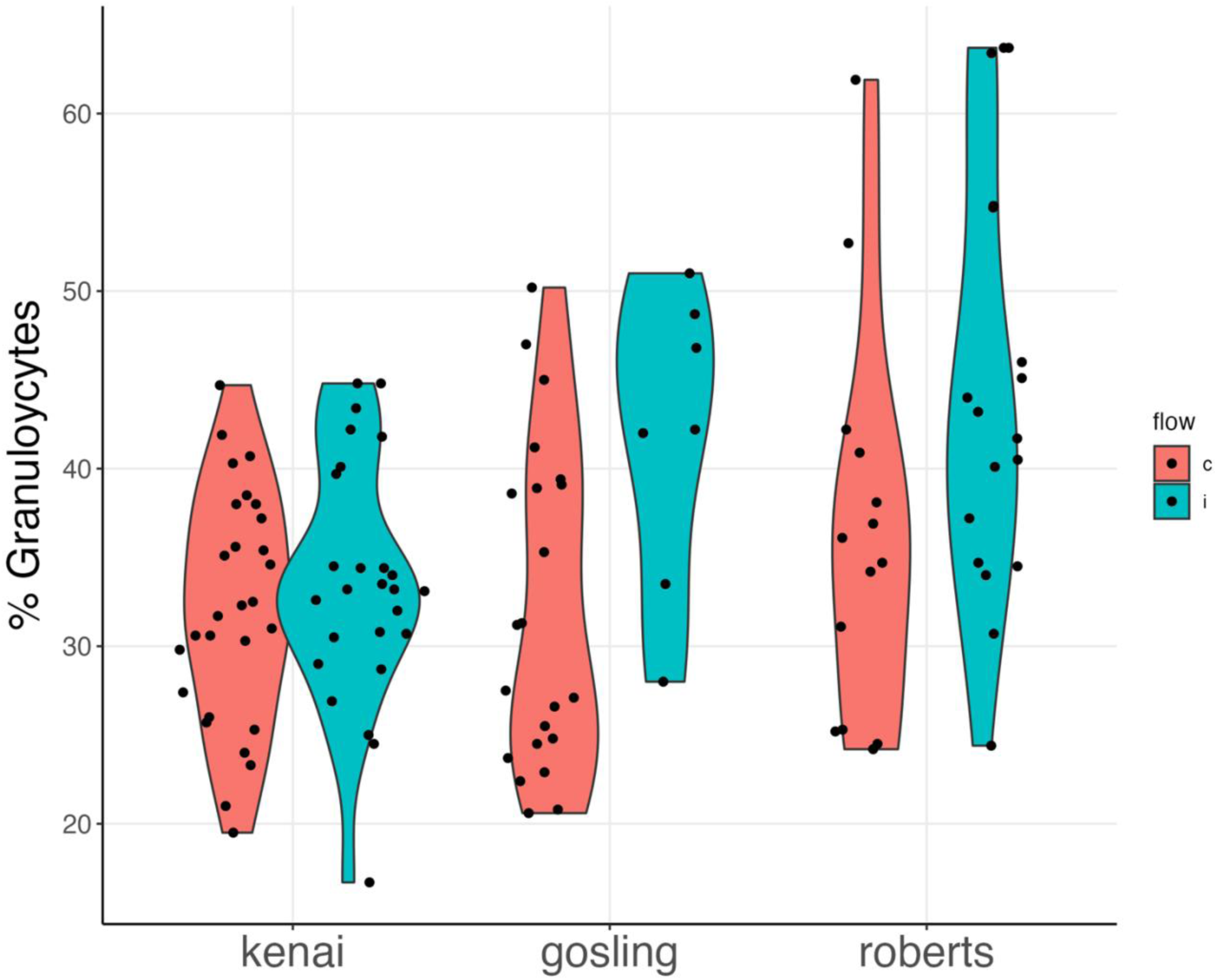
Percent granulocyte cells in head kidney by population. Color indicates final flow regime: continuous flow (“c” = red) and intermittent flow (“i” = blue).

### Liver and Spleen Mass

The best-supported model for liver mass included population, sex, and flow treatment as fixed effects, and age and family as random effects. Movement between flow environments was not associated with changes in liver size. Fish from Gosling Lake had significantly smaller livers than Roberts Lake fish after controlling for mass (-0.44 vs 0.27 mass-scaled liver size, Cohen’s d = -2.1, t-test p<0.00001, Figure 3). The marine population was intermediate in liver size and not significantly different from the other populations. Livers from fish in the c treatment were significantly larger than those observed in fish from the ci (0.16 vs -0.12, d=0.81, p=0.03) and ic treatments (0.16 vs -0.13, d=0.83, p=0.02), and were also marginally larger than livers sampled from fish in the ii treatment (0.16 vs -0.08, d=0.67, p=0.06). Males averaged smaller livers than females, but this difference was not significant (-0.04 vs 0.04, d = 0.23, p=0.16). There was a positive correlation between fish age and mass-corrected liver size (Pearson’s correlation coefficient=0.50, p<0.00001). Mass-corrected liver size was positively correlated with ROS residMFI (r=0.31, p=0.005), but not granulocyte:lymphocyte ratio (p=0.18).

**Figure 3.**
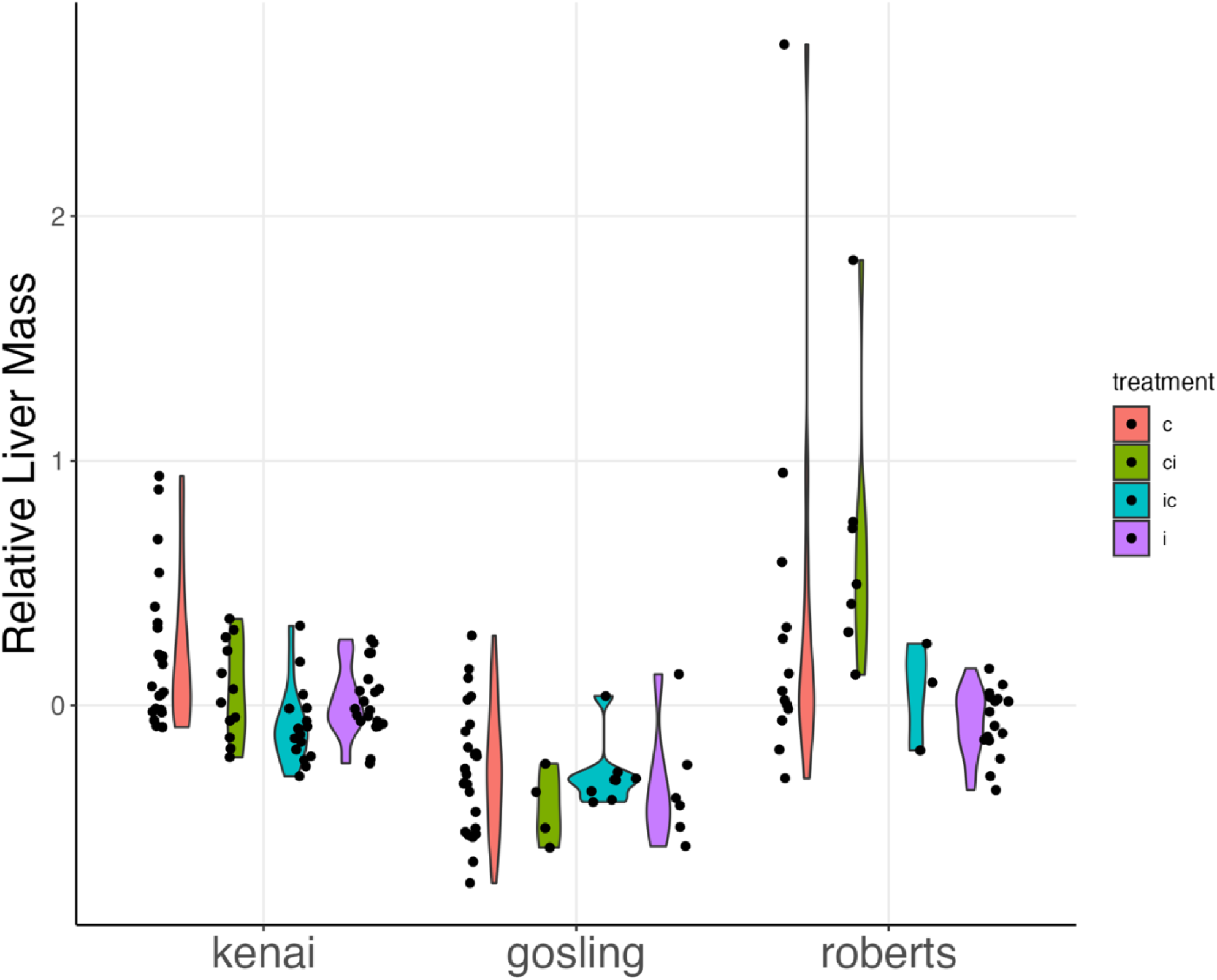
Relative liver mass (corrected for body mass) by population. Color indicates experimental treatment: continuous only (“c” = red), continuous to intermittent (“ci” = green), intermittent to continuous (“ic” = blue), and intermittent only (“i” = purple).

**Figure 4:**
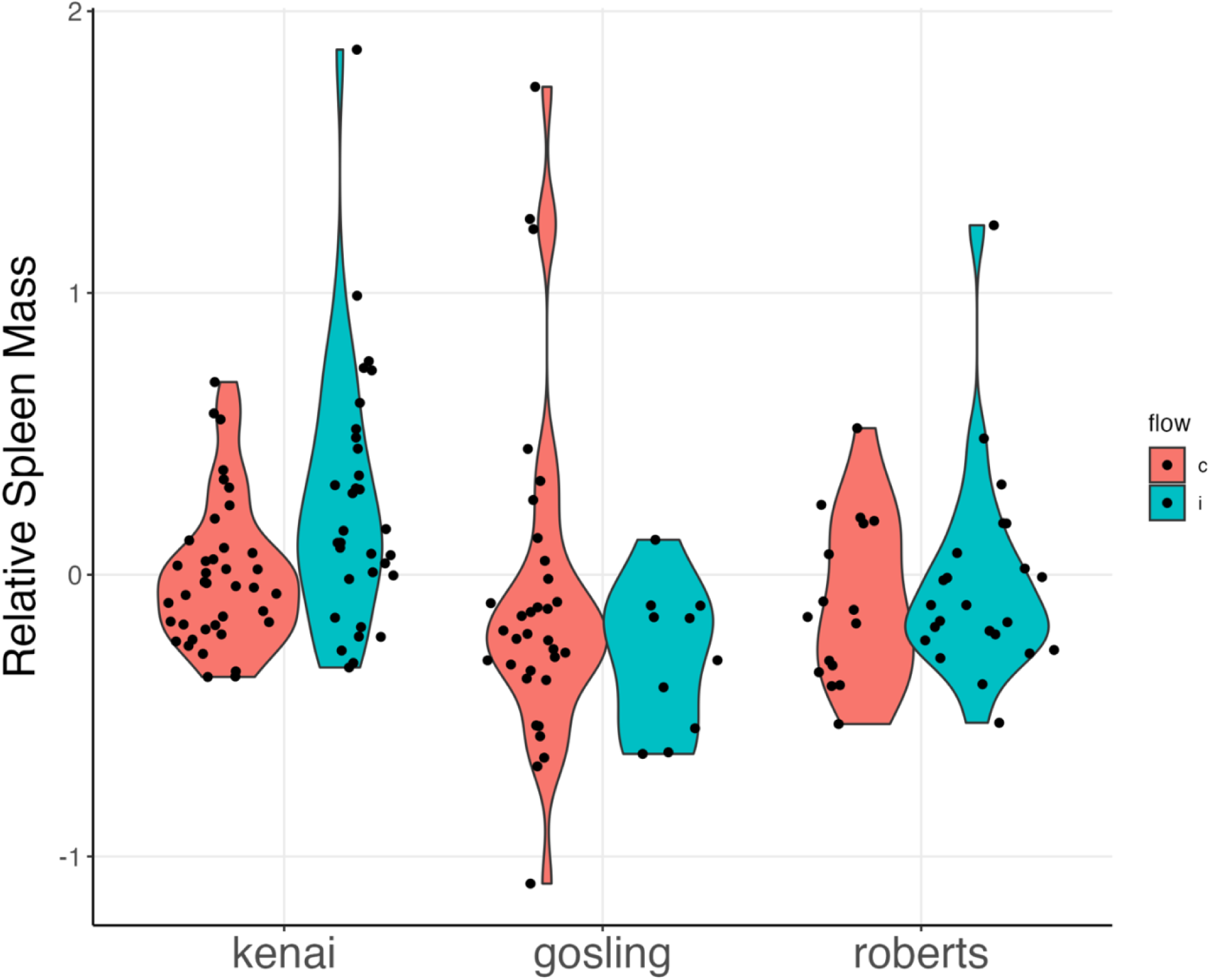
relative spleen mass (corrected for body mass) by population. Color indicates final flow regime (red – continuous flow, blue – intermittent flow)

The best-supported model for spleen mass included fixed effects of population, sex, and final flow environment and age and family as random effects. The marine fish had larger spleens than individuals from both Roberts Lake (0.16 vs -0.08 mass corrected spleen size, d = 0.49, p = 0.01, Figure 5) and Gosling Lake (0.16 vs -0.08, d = 0.55, p = 0.01). Male fish also had larger spleens than females (0.11 vs -0.11, d=0.45, p=0.01), and fish in intermittent flow had marginally larger spleens than continuous flow (0.06 vs -0.07, d = -0.28, p=0.05). Movement between flow environments did not affect spleen mass. There was no significant correlation between mass-corrected spleen size and fish age, ROS production, or granulocyte:lymphocyte ratio (Pearson’s correlation coefficient, p>0.1).

### Tank Microbe Communities

Despite collecting and submitting 20 microbial samples for 16S library preparation and sequencing, we only recovered usable data from four tanks: three continuous flow and one intermittent. We were unable to acquire additional tank microbial samples because the experiment had concluded by the time that we learned of the sequencing error (i.e., we no longer had access to the relevant fish populations in each tank environment). Although the small number of tanks limits our confidence in measurements of microbial differences between water flow environments, we nonetheless identified several patterns. 2D ordination (Figure 1) separated the filter and swab samples along the first NMDS axis, and the intermittent flow tank separated from the three continuous flow tanks along the second axis. The intermittent flow tank also had the highest Shannon diversity index score (Figure S1), though it did not diverge greatly from the continuous flow tanks. We generated a list of the top ten ASVs that had the largest difference in read counts between the tank environments. ASVs in the genera *Sediminibacterium*, *Polynucleobacter*, and *Undibacterium* were overrepresented in both filter and swab samples in the intermittent tank, whereas an ASV in the genus *Sphingopyxis* was underrepresented in the intermittent samples compared to continuous flow.

## Discussion

We leveraged evolved differences in immunity across stickleback populations to investigate how variation in the laboratory environment affects immune function. We found that simply transferring fish between tanks with constant or intermittent water flow, a relatively modest environmental change, led to significant changes in oxidative burst responses and composition of head kidney cells. Housing fish in tanks with intermittent water flow not only drove significantly higher levels of ROS production in granulocytes but also increased the proportion of granulocytes in HK tissue. Notably, most granulocytes in stickleback HKs are neutrophils that drive ROS production during oxidative burst responses (Fuess and Bolnick 2023). Eighteen weeks after fish were transferred into new water flow conditions, the magnitude of both immune measures approximated levels observed in fish that had been housed in each environment for a long period. This demonstrates that environmentally induced flexibility in stickleback innate immune responses is a relatively rapid process.

Apart from classic measures of innate immunity, we also tested the impacts of water flow regime on liver and spleen size. After controlling for random family effects, fish that ended the experiment in intermittent flow tanks had significantly higher SSI, consistent with a plastic shift to greater innate immune allocation. In contrast, there was no simple relationship between HSI and water flow treatment. Stable (non-transplant) rearing under constant water flow yielded significantly higher HSI than the other treatment groups. We had too few tanks available to perform sham transfers (i.e., c-c or i-i), so we cannot rule out a handling effect in the ci or ic treatments. However, this seems unlikely given that fish were allowed to acclimate to their new tanks for approximately eighteen weeks, and handling effects on stickleback immune status are most pronounced within the first two weeks after manipulation (Henrich et al. 2014). Gosling Lake fish had the lowest HSI overall. This was likely due to this population’s fast growth rate, which is also associated with liver size in fish (Pelletier et al. 1994). Marine and Roberts Lake fish did not differ in relative liver size even though marine fish are significantly larger.

All three fish populations displayed a similar magnitude of flexibility for ROS production, granulocyte proportion, and SSI, but population of origin was by far the strongest predictor of overall immune metrics (Table 1). When considering the variance explained by respective best fit models (calculated as total sum of squares), population accounted for 89% of the variance in ROS production, 42% of the variance in granulocyte proportion, and 43% of the variance in SSI. These results closely match previous research describing the evolution of heritable immune differences between stickleback in Gosling and Roberts Lakes (Weber, Steinel, et al. 2017; Weber et al. 2022). Specifically, all studies observed a large difference in oxidative burst capacity (i.e., the Roberts Lake population consistently mounts much larger responses than Gosling), as well comparable differences in granulocyte proportions. The main difference with previous work is that we observed lower absolute ROS levels in the present study. It is unclear whether this absolute difference reflects immunological changes in the natural populations, environmental differences in aquaria habitats, or measurement variation arising from using different flow cytometry platforms. However, it is notable that the population difference in oxidative burst response between Gosling and Roberts Lake has persisted despite the large genomic turnover following the introgression event in Gosling Lake (Flanagan et al. 2025). Specifically, the Gosling Lake fish used in this study were sampled at a timepoint (i.e., 2019) when most animals in that population had evolved a pro-fibrosis response to tapeworm infections. Prior to the introgression event Gosling Lake fish failed to produce fibrosis upon infection (Weber et al. 2017, Weber et al 2022). Previous work also found a significant positive association between fibrosis and oxidative burst capacity in F2-hyrbids between Gosling and Roberts Lake fish (Weber et al. 2022), suggesting that the two traits are functionally linked. The persistently low levels of ROS-production in post-introgression, fibrosis-producing Gosling fish suggest that the mechanistic connection between these traits merits further investigation.

**Table 1:** coefficients from top-ranked model of each trait of interest. 95% confidence interval in parenthesis.

| Trait | Predictor | Beta | SE | p | R<sup>2</sup> |
| --- | --- | --- | --- | --- | --- |
| residROS | intercept | -0.06 (-0.38,0.24) | 0.14 | 0.63 | (0.61,0.70) |
|  | pop-Gosling | -0.53 (-0.88,-0.18) | 0.15 | 0.01 |  |
|  | pop-Roberts | 0.50 (0.14,0.85) | 0.15 | 0.01 |  |
|  | treatment-ci | 0.22 (0.05,0.39) | 0.09 | 0.01 |  |
|  | treatment-ic | 0.18 (-0.01,0.38) | 0.10 | 0.06 |  |
|  | treatment-i | 0.07 (-0.09,0.25) | 0.09 | 0.38 |  |
| % Gran | intercept | -0.19 (-0.69,0.29) | 0.19 | 0.35 | (0.18,0.39) |
|  | pop-Gosling | 0.09 (-0.47,0.66) | 0.23 | 0.70 |  |
|  | pop-Roberts | 0.44 (-0.12,1.01) | 0.22 | 0.10 |  |
|  | sex-m | -0.1 (-0.27,0.05) | 0.08 | 0.19 |  |
|  | flow-int | 0.21 (0.03,0.38) | 0.09 | 0.02 |  |
| Liver mass | intercept | 0.33 (-0.06,0.73) | 0.16 | 0.09 | (0.53,0.70) |
|  | fish mass | 0.95 (0.77,1.12) | 0.09 | 0.00 |  |
|  | pop-Gosling | -0.7 (-1.17,-0.23) | 0.20 | 0.01 |  |
|  | pop-Roberts | 0.1 (-0.36,0.56) | 0.19 | 0.61 |  |
|  | sex-m | -0.11 (-0.21,-0.02) | 0.05 | 0.02 |  |
|  | treatment-ci | -0.18 (-0.34,-0.02) | 0.08 | 0.02 |  |
|  | treatment-ic | -0.21 (-0.36,-0.06) | 0.08 | 0.01 |  |
|  | treatment-i | -0.15 (-0.29,-0.01) | 0.07 | 0.03 |  |
| Spleen mass | intercept | 0.07 (-0.06,0.20) | 0.07 | 0.29 | (0.40,0.40) |
|  | fish mass | 0.8 (0.62,0.98) | 0.09 | 0.00 |  |
|  | pop-Gosling | -0.43 (-0.64,-0.22) | 0.11 | 0.00 |  |
|  | pop-Roberts | -0.24 (-0.4,-0.08) | 0.08 | 0.00 |  |
|  | sex-m | 0.13 (0.00,0.26) | 0.07 | 0.04 |  |
|  | flow-int | 0.13 (0.00,0.26) | 0.07 | 0.05 |  |

Although we previously found that marine and freshwater stickleback differ in their susceptibility to a tapeworm parasite (Weber, Kalbe, et al. 2017), to our knowledge this is the first study to assess oxidative burst capacity in marine stickleback. Interestingly, the intermediate ROS levels in the ancestral marine fish are consistent with a QTL experiment using F2-hybrids between Gosling Lake and Roberts Lake fish, which identified several chromosomal loci that either up- or down-regulate ROS production (Weber et al. 2022). While several other studies have highlighted the utility of population level comparisons of immune variation (Robertson et al. 2016; Peuß et al. 2020), this remains a relatively neglected component of most experimental designs (Sasser and Weber 2023).

While tanks in this experiment shared a common, UV-filtered water source, and all fish were fed the same diet, we hypothesized that several features of the intermittent flow treatment would favor increased microbial diversity. The larger volume of water and much lower rate of water changes should provide greater opportunity for microbes to accumulate. The intermittent flow tanks also housed bubbling sponge biofilters. These are designed to accumulate denitrifying bacteria that remove ammonia waste secreted from fish, which can accumulate at toxic levels in low-flow environments. Our results, with the caveat that they represent a very low number of replicate tanks, support our microbial prediction. The intermittent flow tank had the highest level of microbial diversity in swabs taken from the tank outflow pipes, and the second highest diversity from filtered water samples (Figure S1). We also observed a divergence in microbe communities when accounting for ASV abundance. Whether considering microbes filtered directly from the water or from a swab of the tank outflow, 16S divergence along the first NMDS axis was strongly captured by flow regime (Figure S2). If confirmed when using larger sample sizes, finding a repeatable association between water flow environment and microbial variation would beg several questions: Do changes in tank microbial environment alter the fish microbiome and, if so, are the effects general or specific to areas such as the skin or gut? And are shifts in host immunity caused by microbial exposure, alterations in microbiome, or both events? More directed experiments are needed to address these questions, but our results do at least support a microbe-immune connection. The higher ROS production and granulocyte proportions of HK cells from stickleback in intermittent flow tanks suggests an increased systemic immune stress (Speirs et al. 2024). This is consistent with a bacterial response, potentially mediated via migration of antigen presenting cells from the periphery to the HK (Iliev et al. 2013).

## Data availability

All data are available in a private repository on Dryad: http://datadryad.org/share/LINK_NOT_FOR_PUBLICATION/wm1ejtowfvcgNbRRTcaX2rcWwxtq_6PfVUB4Irk403I. A publicly accessible link will be made available once the manuscript is approved for publication.

---

**Supplemental Table 1:** sample size by population, family, and treatment.

| Population | Family | Treatment | Sample size |
| --- | --- | --- | --- |
| Gosling Lake | Gos14 | C | 7 |
|  | Gos18 | CI | 1 |
|  |  | C | 5 |
|  | Gos35 | C | 5 |
|  | Gos36 | C | 3 |
|  | Gos37 | CI | 5 |
|  |  | IC | 1 |
|  |  | I | 5 |
|  |  | C | 3 |
|  | Gos38 | IC | 8 |
|  |  | I | 1 |
| Kenai River Estuary (marine) | KK1 | I | 10 |
|  |  | IC | 10 |
|  |  | C | 10 |
|  | KK5 | I | 10 |
|  |  | IC | 7 |
|  |  | CI | 12 |
|  |  | C | 12 |
| Roberts Lake | RR1 | C | 4 |
|  | RR2 | CI | 4 |
|  |  | C | 4 |
|  | RR28 | I | 10 |
|  | RR29 | IC | 3 |
|  |  | I | 6 |
|  | RR30 | CI | 4 |
|  | RR31 | C | 5 |

**Supplemental Figure 1:**
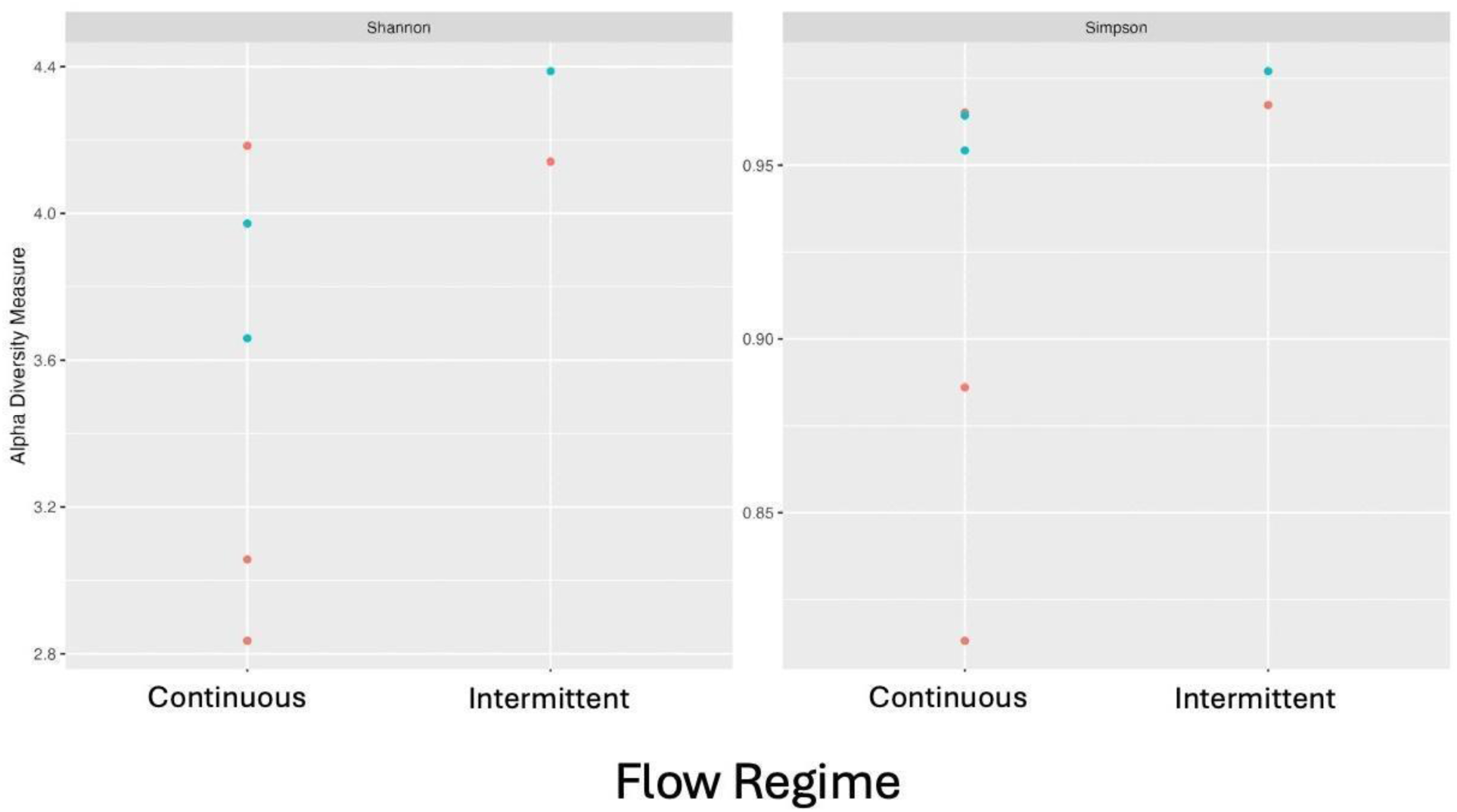
diversity indices of tank microbial communities based on 16S data by flow regime. Color indicates sampling location (blue – tank outlet swab, red –tank water sample).

**Supplemental Figure 2:**
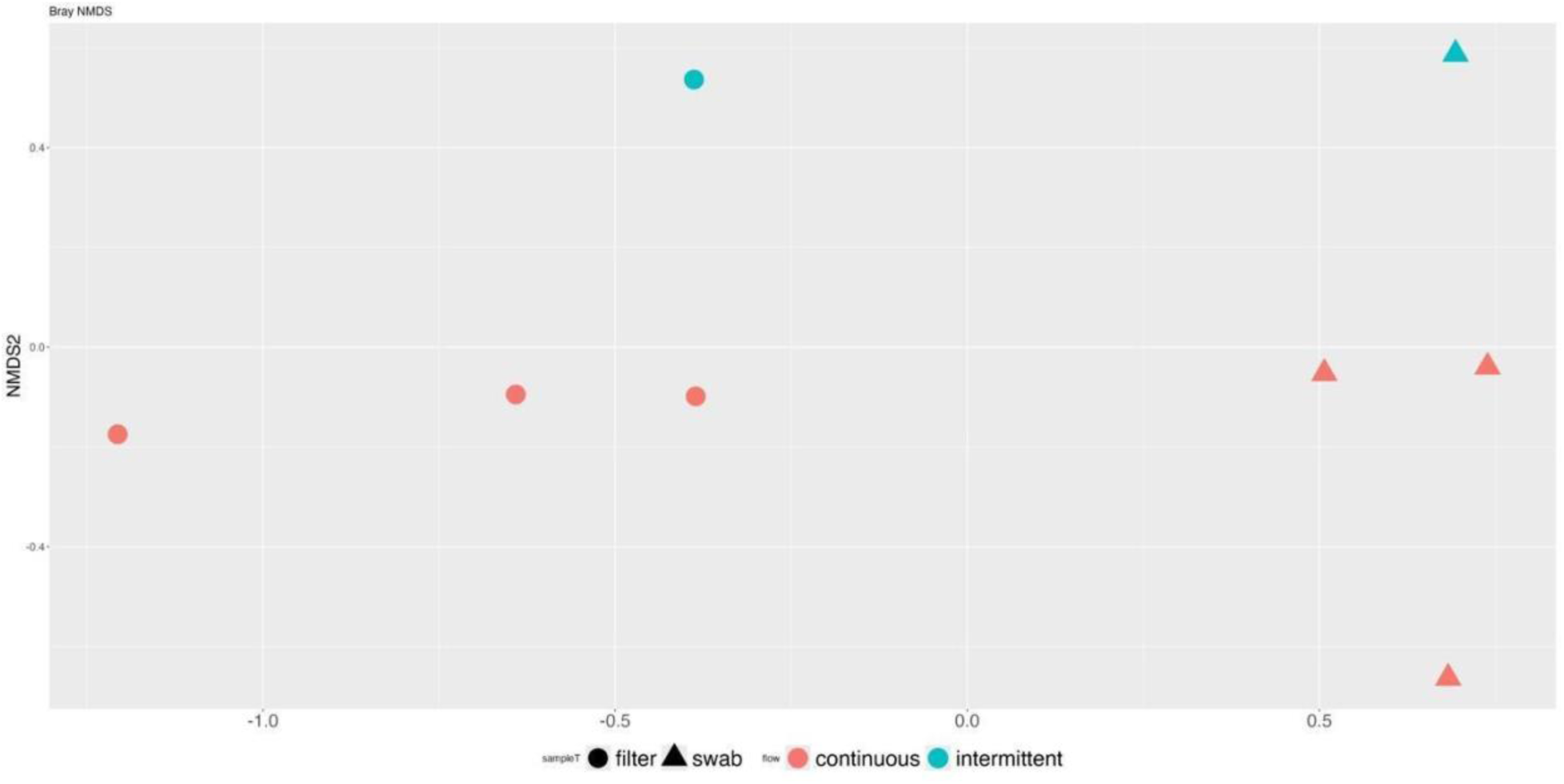
Bray-Curtis non-metric multidimensional ordination scaling plot of tank microbial communities based on 16S data. Color indicates flow regime (red – continuous flow, blue – intermittent flow) and shape indicates final sampling location (square – tank outlet swab, triangle – tank water sample).

